# Same, Same, but Different: Molecular Analyses of *Streptococcus pneumoniae* Immune Evasion Proteins Identifies new Domains and Reveals Structural Differences between PspC and Hic Variants

**DOI:** 10.1101/2020.08.11.246066

**Authors:** Shanshan Du, Claudia Vilhena, Samantha King, Alfredo Sahagun, Sven Hammerschmidt, Christine Skerka, Peter F. Zipfel

## Abstract

PspC and Hic proteins of *Streptococcus pneumoniae* are some of the most variable microbial immune evasion proteins identified to date. Due to structural similarities and conserved binding profiles it was assumed over a long time that these pneumococcal surface proteins represent a protein family, comprising eleven subgroups. Recently, however, by evaluating more proteins larger diversity of individual proteins became apparent. In contrast to previous assumptions a pattern evaluation of six PspC and five Hic variants, each representing one of the previously defined subgroups, revealed distinct structural and likely functionally regions of the proteins, and identified nine new domains and new domain alternates. Several domains are unique to PspC and Hic variants, while other domains are shared with other *S. pneumoniae* and bacterial virulent determinants. This understanding improved pattern evaluation on the level of full-length proteins, allowed a sequence comparison on the domain level and furthermore identified domains with a modular composition. This novel concept allows a better characterization of variability, and modular domain composition of individual proteins, enables a structural and functional characterization at the domain level and furthermore shows substantial structural differences between PspC and Hic proteins. Such knowledge will also be useful for molecular strain typing, characterizing PspC and Hic proteins from new clinical *S. pneumoniae* strains, including those derived from patients who present with pneumococcal hemolytic uremic syndrome. Furthermore this analysis explains the role of multifaceted intact PspC and Hic proteins in pathogen host interactions. and can provide a basis for rational vaccine design.

**Author Summary:** The human pathobiont *Streptococcus pneumoniae* expresses highly polymorphic PspC or Hic proteins, which bind a repertoire of host immune regulators and combine antigenic variation with conserved immune evasion features. Understanding domain composition of each protein encoded by more than 60 000 *pspC* or *hic* genes deposited in the data banks defines their diversity, a role in immune escape and can furthermore delineate structure function approach for single protein domains. PspC and Hic proteins show variable domain composition and sequence diversity, which explain differences in binding of human regulators and likely in immune escape. The results of our analyses provide insights in the domain composition of these diverse immune evasion proteins, identifies new domains, defines domains which are unique to PspC or Hic variants, and identifies domains which are shared with other bacterial immune evasion proteins. These data have implication on cell wall attachment, surface distribution and in immune escape.

## Introduction

### The pathobiont Streptococcus pneumoniae

*Streptococcus pneumoniae* (the pneumococcus) is the major cause of community-acquired pneumonia. In addition, this human pathogenic Gram-positive bacterium can cause otitis media and may also cause acute life-threatening invasive infections such as meningitis and even sepsis (1–4). Malnutrition and *S. pneumoniae* infections are the major cause of childhood mortality worldwide. Pneumonia account for approximately 16 percent of the 5.6 million of deaths among children under five years old, killing around 808,000 children in 2016 according to the United Nations Children’s Fund (UNICEF) and the World Health Organization (WHO)(5–7). At any point in time pneumococci can reside asymptomatically in the upper respiratory tract of about 50% of children, from where they can be transmitted to other persons. Based on differences of the polysaccharide capsule so far 97 serotypes are identified (8).

Pneumococcal diseases are widespread both in developing and developed countries and antibiotic resistant strains are arising. The increase in microbial resistance to antibiotics makes it important to identify new virulence determinants, to understand the diversity of these determinants and also to define the immune escape strategies of this relevant pathogenic bacterium (9–11). In addition, vaccines with higher serotype coverage or serotype-independent vaccines are needed in order to combat the pathogen.

Immune and in particular complement evasion is critical for *S. pneumoniae* and for all the ability of all human pathogenic microbes to cause infections. Common patterns regarding complement evasion and binding of human complement and immune regulators are emerging (12–16). Thus, it is important to understand the exact role of individual pneumococcal virulence determinants, in particular their role of complement evasion, the topology of the capsule and surface location of virulence determinants (17–19).

### PspC and Hic represent related *S. pneumoniae* surface proteins

PspC and Hic proteins are important pneumococcal immune evasion proteins and adhesins and represent promising vaccine candidates (20). The majority of virulent *S. pneumoniae* strains express at least one Psp or Hic variant, and strains that have the *pspC/hic* genes deleted show significant amelioration of lung infection in mice, nasopharyngeal colonization, and bacteremia (21).

### PspC and Hic proteins as central pneumococcal immune evasion proteins

Initially, PspC was identified as an adhesin, which targets the secretory component of the secreted Immunoglobulin A (sIgA) and polymeric IgA receptor (pIgR)(22). Because *pspC* and *hic* genes were identified independently by several groups, different names were originally given, including CbpA (choline-binding protein A), SpsA (secretory IgA binding protein), PbcA (C3-binding protein A), or Hic (Factor H binding inhibitor of complement)(23–33).

PspC and Hic proteins are attached to the bacterial cell wall, and are surface-exposed adhesins and immune evasion proteins. PspC proteins with their C-terminal choline-binding anchors attach non-covalently to the phosphorylcoline (PCho) moiety of the teichoic acids (TAs) and Hic proteins, displaying a C-terminal LPsTG motif, are covalently linked to the peptidoglycan via the sortase A. This suggests that proteins attach with different strengths and likely also to a distinct depth in the cell wall, which C-terminal region suggests that the preceding part of the protein spans the capsular polysaccharides and that the N-terminal part is extending beyond the capsule.

As central immune evasion proteins, PspC and Hic proteins bind several human plasma proteins including Factor H, C3, C4BP, Plasminogen, thrombospondin-1, and vitronectin (22–36). These multifunctional proteins represent one of the most diverse group of immune evasion and adhesive proteins recognized to date (35, 36). PspC and Hic proteins have *a* mosaic structure, comprising distinct regions, consisting of multiple domains. Furthermore a substantial overlap of domains exist between PspC and Hic variants. Standard domain or sequence-based comparison among members of this protein family is complex due to structural differences and variable domain composition. Currently, the protein NCBI databank lists 54909 entries for PspC or Hic and 11817 entries for CbpA, encoding both full-length proteins and partial protein sequences (march 06, 2020; NCBI www.ncbi.nlm.nih.gov/protein). The individual entries show homology, but also exhibit considerable variations in structure and sequence. Single PspC and Hic proteins show variable domain patterns, different variants of these proteins combine domains in different ways, and apparently not all domains are identified so far.

#### Mosaic-structured PspC and Hic proteins

Our understanding of these important pneumococcal immune evasion proteins is currently still fragmented. Thus, defining the exact domain composition of individual PspC and Hic variants, or to correlate phenotypes with disease forms, is important for better understanding the role of each protein, for structural predictions, for localizing binding sites of host ligands, for understanding precise domain function(s), and for characterizing strain specific differences.

Based on overall sequence similarities PspC and Hic variants were initially considered to belong to one group of pneumococcal immune evasion proteins. Initial analyses by Brooks Walter in 1999 and Iannelli et al. in 2001 revealed both sequence similarity and diversity among PspC and Hic proteins (37, 38). Ianelli et al. identified several domains for the evaluated 43 PspC and Hic proteins including the leader peptide, α-helical regions with a seven-amino acid periodicity, repeat domains, a proline-rich stretch followed by either a choline-binding or sortase-dependent anchor (38). At that time, the cell wall anchors were used as criterion to differentiate between PspC and Hic family proteins and based on sequence differences six PspC-type and five Hic-type clusters were defined. However, still today no precise criteria exist regarding cluster specific domain composition or domain characteristics. Because the domain patterns as well as borders of single domains are not well-defined, a straightforward variant designation e.g. of existing but also of newly identified *pspC and hic* genes, or genes from novel clinical pneumococcal isolates, is difficult or even impossible (39).

#### Aim of the study

Thus far, the internal or external position of each domain remains unclear, as do the precise borders of the regions and of the domains. We do not exactly know which domain(s) are indeed integrated into the bacterial cell wall, which domain(s) span the capsule, or which domains are externally positioned. Given these limitations, and the heterogeneity among the proteins, we aimed to evaluate the structure and domain composition of six PspC and five Hic variants, each representing one of the clusters defined by Ianelli *et al*. We further aimed to obtain evidence on domain composition and position. By evaluating the domain pattern of each variant, we identified nine new domains, illustrate structural as well as ositional differences between the full length proteins, between N and C-terminal s, and between PspC and Hic proteins. Furthermore this comparison also identified subvariants on the domain level.

## Results

### Global Similarity of PspC and Hic Variant Proteins

#### Selection of PspC and Hic variants

One protein from each variant cluster as defined by Ianelli et al. was selected (38). The variants included the six PspC variants, i.e. PspC1.1, PspC2.2, PspC3.1, PspC4.2, PspC5.1, PspC6.1, and five Hic variants, Hic/PspC7.1, Hic/PspC8.1, Hic/PspC9.1, Hic/PspC10.1, Hic/PspC11.1. At the date of the cluster designation Ianelli *et al.* considered the PspC and Hic variants as one protein family and used a PspC nomenclature for both protein groups (38). To appreciate the Hic type character and at the same time follow the nomenclature suggested by Ianelli *et al* we combine the Hic and PspC designations **(Figure 1A)**. The selected proteins vary in size and mass, with PspC1.1 as the largest protein having a length of 929 aa and a molecular mass of 110 kDa, while Hic/PspC8.1 is the smallest protein with a length of 503 aa and a mass of 65 kDa (Supplementary Table I). When compared to the well-characterized PspC3.1 protein (strain D39), the overall sequence amino acid identity of the six PspC proteins ranged from 51 to 82%. In contrast, the five Hic variants showed a less pronounced identity which ranged from 15 to 26%. Thus suggesting functional differences between the PspC and Hic variants **(Figure 1B).**

**Figure 1:**
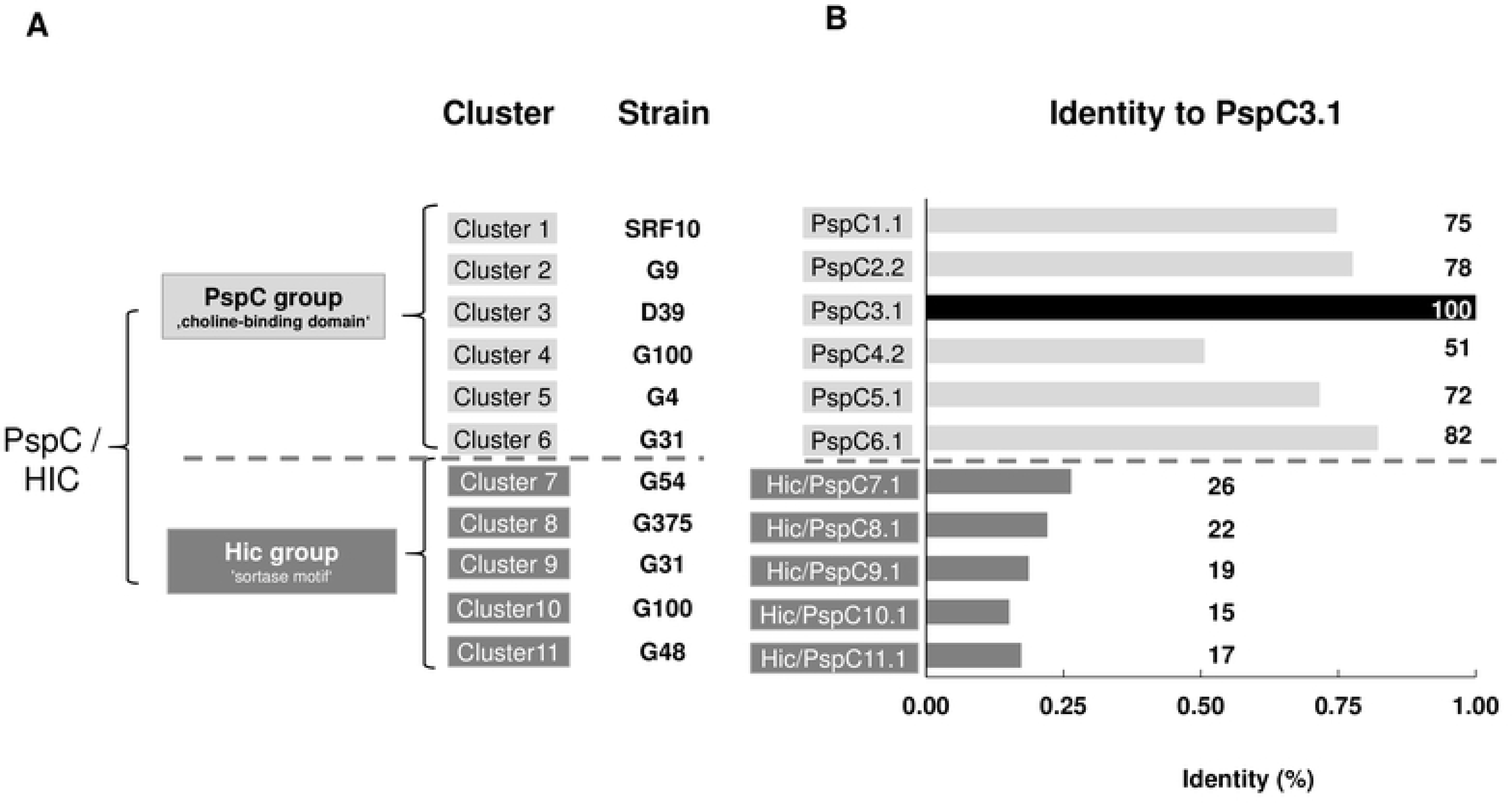
Diversity of PspC and Hic cluster variants. PspC and Hic proteins were initially considered to represent one protein class that based on the different surface anchors can be divided into two major clusters. **A:** PspC variants with choline-binding domains representing the PspC group, and Hic variants having sortase LPsTG motifs for cell wall anchoring the second namely Hic group. For each group additional cluster or subgroups were identified. For the analysis one variant from each cluster was selected, i.e. for PspC group: PspC1.1, PspC2.2, PspC3.1, PspC4.2, PspC5.1, PspC6.1; and for the Hic group: Hic/PspC7.1, Hic/PspC8.1, Hic/PspC9.1, Hic/PspC10.1, Hic/PspC11.1. **B: Overall sequence homology among the selected cluster variants.** Amino acid homology of the indicated full-length protein variants was compared to PspC3.1, which was used as reference. The sequence variation shows differences for the six selected PspC and the five Hic variants. This difference is indicative for compositional variation among the two major protein groups.

**Table I:** Human Regulators binding to PspC and Hic variants. Binding of human plasma regulators to PspC and Hic proteins. The binding sites for Factor H has been mapped within the Hypervariable Domain of PspC3.1 and that of sIgA and the extracellular domain of pIgR to the RNYPT motif of Repeat Domains I and II. Binding of C3, C4BP, Plasminogen, Thrombospondin 1, vitronectin have been show to intact *S. pneumoniae* and to full length PspC and Hic proteins, but their binding sites have not been mapped to single domains so far. Interaction of Lactoferrin and IgA is proposed based on homology between PspC and Hic variants with the *S. pneumoniae* immune escape protein PspA, and the homology to the sIgA binding protein of *S. agalactiae*.

| # | Region |  |  | Domain | Sub domains | Class | n | Module | Structure |  | Comment<br>Host Ligand |
| --- | --- | --- | --- | --- | --- | --- | --- | --- | --- | --- | --- |
| 1 |  |  | known | SP |  |  | 11 |  |  |  |  |
| 2 | N-term | | known | HVD | HVD-A, HVD-B,<br>HVD-C | | 11 | | $\alpha$ helix | PspC/Hic<br>specific | Factor H |
| 3 | | | known | RD | RD-I, RD-II | | 7 | | $\alpha$ helix | | sIgA/pIgR |
| | | | | | RDII | PspC | 5 | | $\alpha$ helix | | |
| 4 | | | known | RCD | | | 8 | | $\alpha$ helix | | |
| 5 |  | 1 | new | S<sub>n</sub>D/GS<sub>2</sub> | 3 Positions |  | 10 |  | coiled coil |  |  |
| 6 | | 2 | new | RCE1 | | PspC | 2 | | $\alpha$ helix | | Lactoferrin |
| 7 |  | 3 | new | RCE2 |  |  | 2 |  | A helix |  |  |
| 8 | | 4 | new | PspA<br>related | | PspC | 2 | | $\alpha$ helix | in PspA | |
| 9 | | 5 | new | R-type | | | 3 | | $\alpha$ helix | | IgA |
| 10 | | 6 | new | EPRD | | | 4 | | $\alpha$ helix | | |
| 11 | | 7 | new | IgA | | | 4 | | $\alpha$ helix | *S. agalactiae* | |
| 12 | | 8 | new | VS4.2 | | PspC | 1 | | $\alpha$ helix | Specific | |
| 13 | | 9 | new | VS11.1 | | Hic | 1 | | $\alpha$ helix | Specific | |
| 14 | C-term |  | known | PRD | PRD-IA,<br>PRD-IB | PspC | 5 | Modular | coiled coil | also in PspA | Cell wall spanning? |
|  |  |  | new |  | PRD-II | PspC | 1 | Modular | coiled coil | ? |  |
|  |  |  | new |  | PRD-III | Hic | 1 | Modular | coiled coil | ? |  |
|  |  |  | new |  | PRD-IV | Hic | 4 | Modular | coiled coil | ? |  |
| 15 | | | known | anchor | CBD | PspC | 6 | Modular | $\beta$ sheets | several | Anchor |
| 16 |  |  | known |  | LPsTG | Hic | 5 |  | coiled coil | many | Anchor |

#### PspC3.1 as a prototype PspC

PspC3.1 was selected as prototype and used for analyzing structure and domain composition. PspC3.1 has a signal peptide that directs the protein to export. The protein has an externally oriented N-terminal region and is integrated into the teichoic acids of the bacterial cell wall via the C-terminal Choline-Binding Domain. Because different regions of these membrane anchored proteins are facing different environments, we hypothesized, that hydrophilic and hydrophobic surroundings, could influence protein structure and composition.

#### Structure and residue composition of PspC3.1

PspC3.1, when evaluated *in silico,* showed three clearly different structural regions. The N-terminal 410 residues form mostly α−helices, followed by a 70 aa long predominately coiled-coil region and a 221 aa long region composed mainly of β-sheets (**Figure 2A).** Given these structural differences the 410 aa long mainly α−helical region was designated as N-terminal region, and the remaining part with the coiled-coil and β-sheet segments and almost lacking α−helices was termed C-terminal region. When the structural regions were aligned with the known domains of PspC3.1, the N-terminal α−helical region included the signal peptide, the Hypervariable Domain, the two Repeat Domains, and the Random Coil Domain. The Hypervariable Domain includes the binding sites for human Factor H and each Repeat Domain includes a binding site for sIgA/polymeric Ig receptor, which is in agreement with an external, orientation. The mostly coiled-coil structured region represents the Proline-Rich Domain (aa 411-482), which is considered a cell wall-spanning and flexible domain and the β-sheet region represented the Choline-Binding Domain (aa 483-701) used for cell wall attachment **(Figure 2B**). The C-terminal Proline-Rich Domain and the Choline-Binding Domain have an inside location (40, 41).

**Figure 2:**
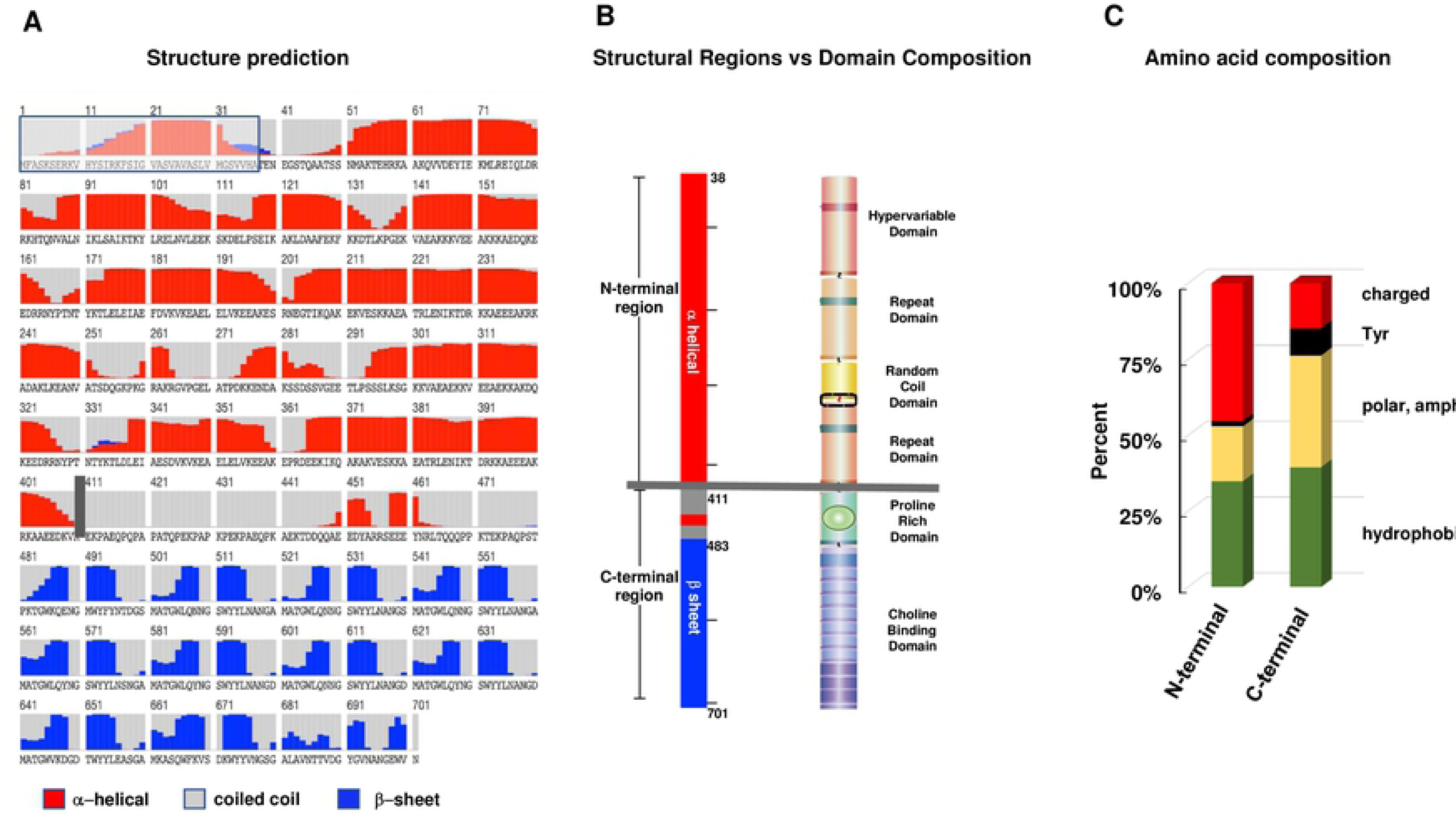
Structural regions and domain position of PspC3.1. *In silico* structure analyses of PspC3.1 dissects distinct structural region. A: The structure of the well-characterized PspC3.1 (strain D39) was evaluated *in silico*. The N-terminal part of the molecule shows a long stretch composed mainly of α-helices (red columns) (aa 1-410), being followed by a 72 aa long coiled-coil structured segment (grey area) and by a 219 aa long segment with β−sheet folds (blue columns). The numbers on top represent the amino acid position within the protein. The signal peptide (positions 1-37) which is cleaved upon processing is shown by the box with grey background and blue lines. The vertical grey bar separating the N-terminal α helical from the coiled coil structured region may represent the position of the bacterial cell wall. **B: Structural regions and domain composition of PspC3.1**. The mainly α−helical structured region (position 38 to 410) is termed the N-terminal region. The remained of the protein that includes the 72 aa coiled-coil structured and the 219 aa mainly β-sheet segment is termed the C-terminal region (left panel). To correlate structural regions with the domain composition, the know domains of PspC3.1 were aligned (right panel). The hypervariable domain, repeat domain I, random coil domain, repeat domain II aligned with the N-terminal, mainly α-helical region. In the C-terminal part the Proline-Rich Domain lined up with the coiled-coil structured region and the Choline-Binding Domain with the β-sheet region. The grey horizontal line separates the N and C-terminal regions and likely marks the border of the cell wall and capsule facing the outside environment. **C: Amino acid composition of N and C-terminal regions.** The amino acid composition was evaluated for each region separately. The N-terminal region is rich in charged residues (48%), has low degree of polar and amphipathic residues (24%), and contains a low fraction of Tyr residues (left panel). The C-terminal region contained a lower fraction of charged residues (22%), had more polar amphipathic amino acids (38%) and more Tyr residues (8%).

#### Amino acid composition

Next we evaluated if the proposed outside and inside environments influence the protein make up. The N-terminal region of PspC3.1 includes 45.3% charged, 18.0% polar and amphipathic residues and has a low fraction of Tyrosines (1,7**%**). The C-terminal region in contrast includes only 15.0% charged residues, has 3an increase percentage or (9.5%) polar and amphipathic amino acids (9.5%) and many Tyrosines (8.9%) (**Figure 2C).** Thus the N-terminal and C-terminal regions of PspC3.1 differ in structure, and amino acid composition.

#### The differences of the N and C-terminal regions are conserved in the PspC and Hic variants

Next we evaluated if the structural composition, as outlined for PspC3.1, is conserved in the other PspC and Hic variants. The N-terminal regions of all analyzed PspC and Hic variants have mostly α-helical structrues, and the C-terminal Proline-Rich Domains have predominantly coiled-coil structures. The PspC specific Choline-Binding Domains have mostly β−sheets, and the Hic specific LPsTG anchors have an α−helical segment following a coiled-coil stretch (**Supplementary Figures 1 and 2**).

In addition the amino acid composition was determined. The N-terminal regions of the six PspC variants contained 35-45% charged residues. In contrast their C-terminal regions contained 16% or less charged residues. The C-terminal domain also included more polar and amphipathic amino acids (32-36%) and were rich in Tyrosine (8.3-9.8%)(**Figure 3A**), The Hic variants contained 28-37% charged residues in their N-terminal regions, and their C-terminal regions had a high fraction of charged (28-41%) and less polar/amphipathic residues (15-21%) than the PspC variants (**Figure 3A)**. Thus, the N and C-terminal regions of the proteins differ in structure and amino acid composition and the C-terminal regions of the PspC and Hic proteins show differences in amino acid composition.

**Figure 3:**
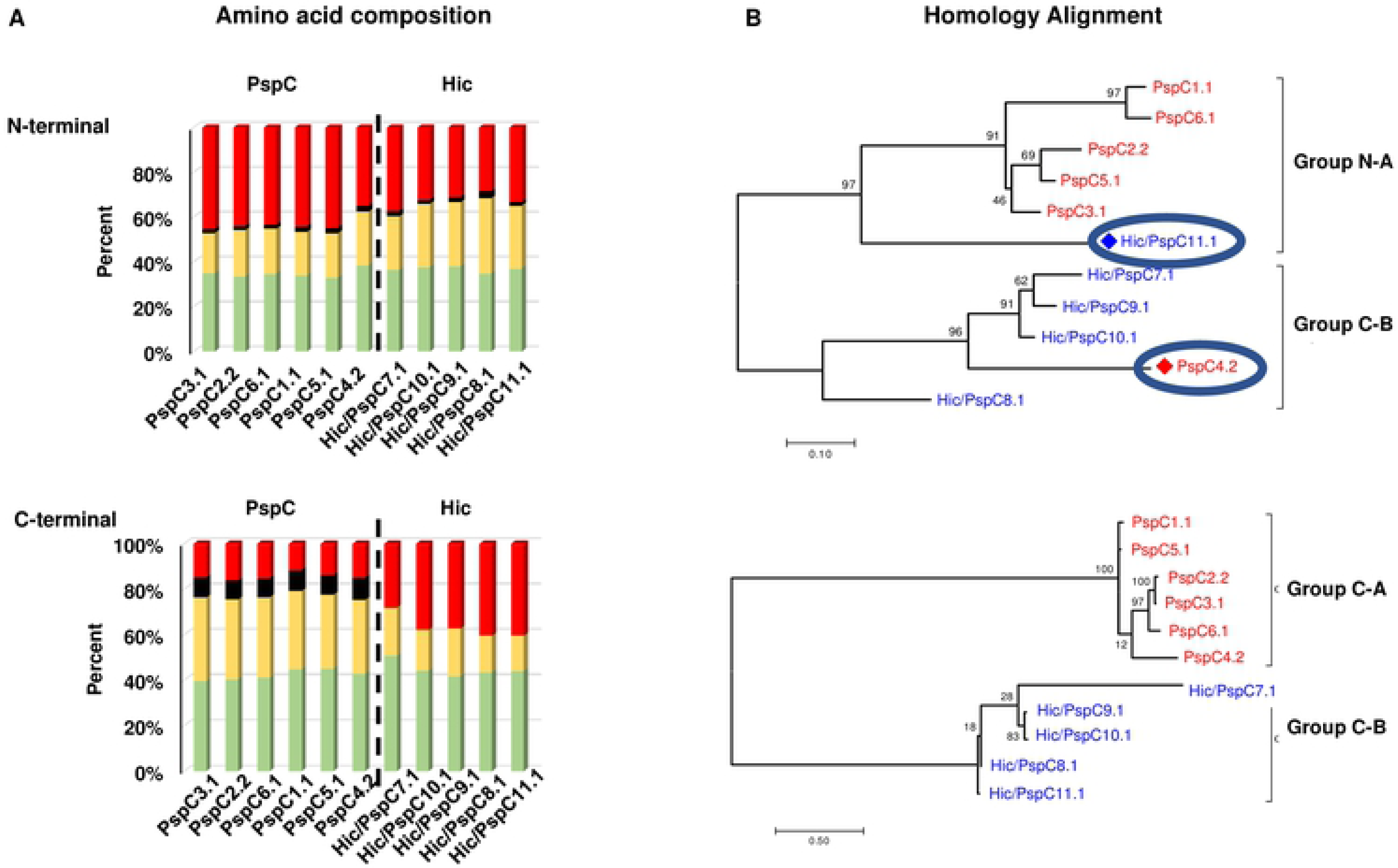
Differences in the N and C-terminal regions of the PspC and Hic variants. A: The N and C-terminal regions of PspC and Hic type proteins differ in amino acid composition. The amino acid composition of the N and C-terminal regions was evaluated for each of the six PspC and the five Hic variants. The N-terminal regions of the PspC and Hic variants are rich in charged residues (35-45%), have a low degree of polar and amphipathic residues, and contain a low percentage of Tyr residues. The PspC variants had also a high portion of charged residues (28-27%)(upper panel). The C-terminal regions of the PspC variants had a lower fraction of charged residues (16 % or less), more polar and hydrophilic residues (32-36% and more Tyr residues (8.3-9.1%). The composition of the C-terminal region of Hic variants differed from that of PspC variants. The C-terminal regions of Hic variants showed more charged residues, a lack of Tyr residues and less polar and amphipathic residues (lower panel). **B: Homology alignment of the N and C- terminal regions of PspC and Hic type proteins.** The homology alignment of the N and C-terminal regions identifies two groups. For the N-terminal regions the first, group A is dominated by PspC type proteins but also includes Hic/PspC11.1. The second N terminal group B is dominated by Hic type proteins but also includes the PspC4.2 variant. The C-terminal regions show a clear separation among the PspC and Hic variants.

The N-terminal regions of the different variants ranged in length from **146** (Hic/PspC8.1) to **633** (PspC5.1) residues. A homology alignment of the N-terminal regions showed two distinct clusters. One N-terminal cluster included five PspC variants (PspC1.1, PspC6.1, PspC2.2, PspC5.1, PspC3.1) and the Hic/PspC11.1 variant while the second N-terminal panel included PspC4.2 and four Hic variants (Hic/PspC7.1, HicPspC9.1, Hic/PspC10.1, Hic/PspC8.1)(**Figure 3B, upper panel).**

The C-terminal regions were more conserved in length, ranging from 236 (PspC5.1) to 348 aa (Hic/PspC8.1) and were clearly separated the PspC and the Hic members. The level of diversity between the C-terminal regions of variants within each group was low indicating that these domains are more highly conserved (**Figure 3B, lower panel).**

#### Domain analyses of PspC and Hic variants

Using PspC3.1 with its five known domains as a blue print, the domain patterns of the other ten cluster variants was evaluated. This approach determined that three domains of PspC3.1, the signal peptide, the N-terminal Hypervariable Domain and Proline-Rich Domains are found in all PspC and Hic variants. All PspC variants use a Choline-Binding Domain, while Hic/PspC proteins have an LPsTG anchor (**Figure 1 and Figure 4**). Repeat Domains and the Random Coil Domain are found mainly in PspC proteins, but not in all variants. Additional sequence stretches were identified in some PspC or Hic variants that did not match with known domains of PspC3.1. These domains were evaluated separately to determine whether counterparts exist in other PspC and Hic variants or whether homologs exist in the protein data bank. This approach identified nine new domains, including one new domain in PspC3.1 and also three new sub variants of the Proline Rich Domain. This extended domain scenario shows that the individual PspC and Hic proteins harbour variable domain numbers, ranging from four (Hic/PspC8.1) to ten domains (PspC4.2**)(Figure 4).**

**Figure 4:**
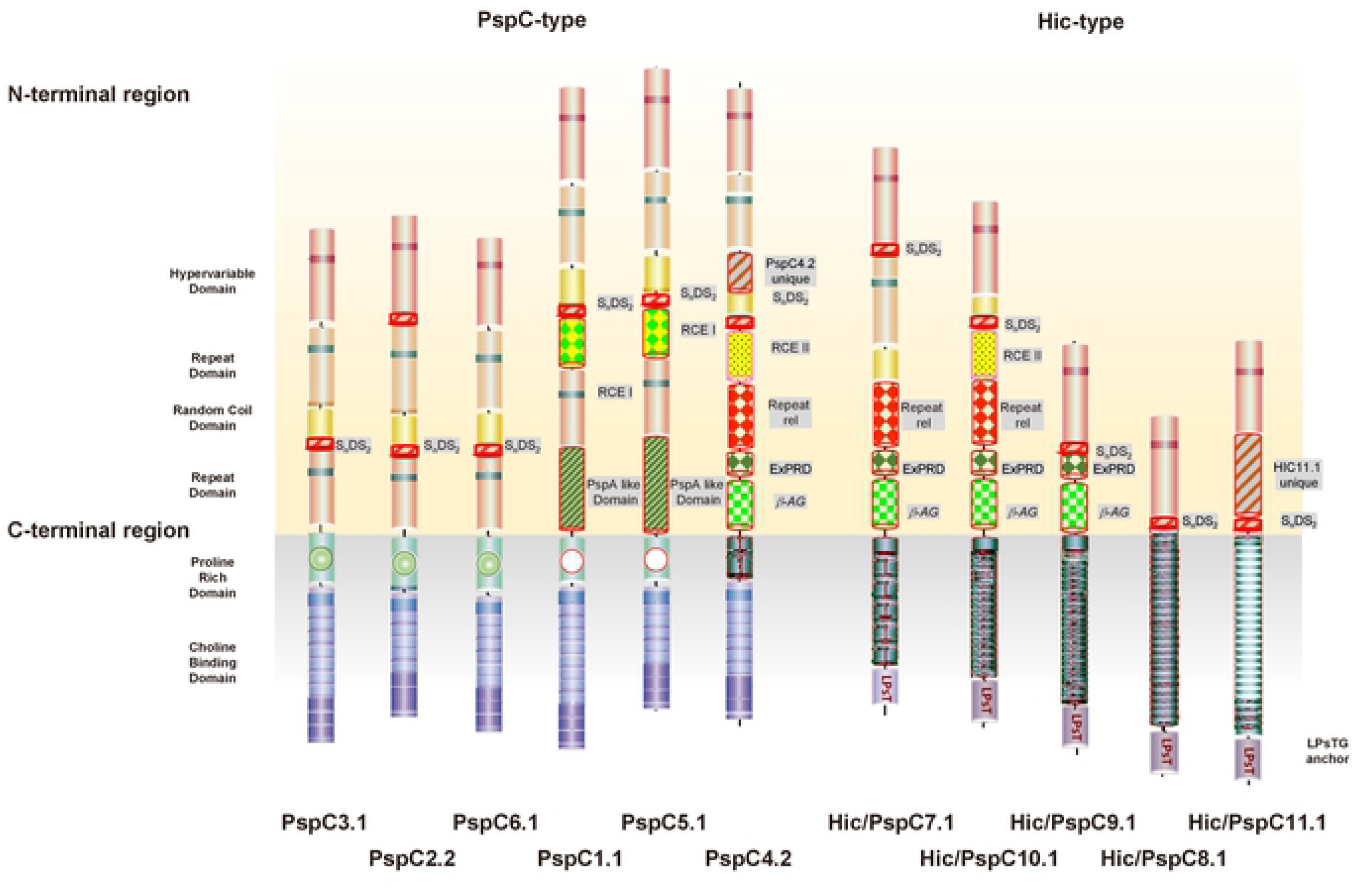
Domain structure of the six PspC and th efive Hic variants. **A: PspC3.1 with the domain architecture** is shown on the left side. The PspC and Hic variants differ in length and in domain number. The proteins each representing one member of the previously identified clusters are arranged based on their overall homology. To reflect the different lengths of the proposed outside and interior regions the proteins are centered along the axis, which separates the N-terminal α-helical region from the C-terminal region. N-terminal regions are shown on yellow and C- terminal regions on a grey background. Proteins are drawn to scale. The signal peptides and the most C-terminal segments of class II proteins, which are cleaved by the transpeptidase sortase are not presented. Know domains as identified for PspC3.1 are shown in filled color. New domains are patterned, and the names are represented on grey background. The mapped binding sites for the human plasma proteins Factor H in the Hypervariable Domain are shown by the purple bar and that of sIgA/pIgR in the Repeat Domains by green bars. Lactoferrin and IgA binding domains are proposed by homology with binding domains of *S. pneumoniae* protein PspA and by the IBC protein from *S. agalactiae*.

#### Known domains of the N-terminal region

The known domains identified in the N-terminal region include:

##### Signal peptide

A highly-conserved 37 aa long N-terminal signal sequence which directs the proteins for export and is cleaved upon processing is present in all PspC and Hic/PspC variants (**Supplementary Figure 3A**).

##### Hypervariable domains

Mature PspC and Hic/PspC proteins expose N-terminal Hypervariable Domains, which are rich in charged residues. The length of the Hypervariable Domains ranged from 91 (PspC4.2) to 113 aa (PspC2.2), and as their name suggests they were highly variable in sequence. Only five residues, **T**_11_,**S**_12_,**I_59_**,**Y_63,_K_96_** (numbering based on PspC3.1) are present in all proteins although additional residues are conserved in several variants. The N-terminal **Hypervariable Domains** appear to be specific for PspC/Hic protein variants (**Supplementary Figure 3B**) and they include a 12 amino acid long region, which in PspC3.1 was identified as Factor H binding region **(Figure 5A**, **Supplementary Figure 3C**).

**Figure 5:**
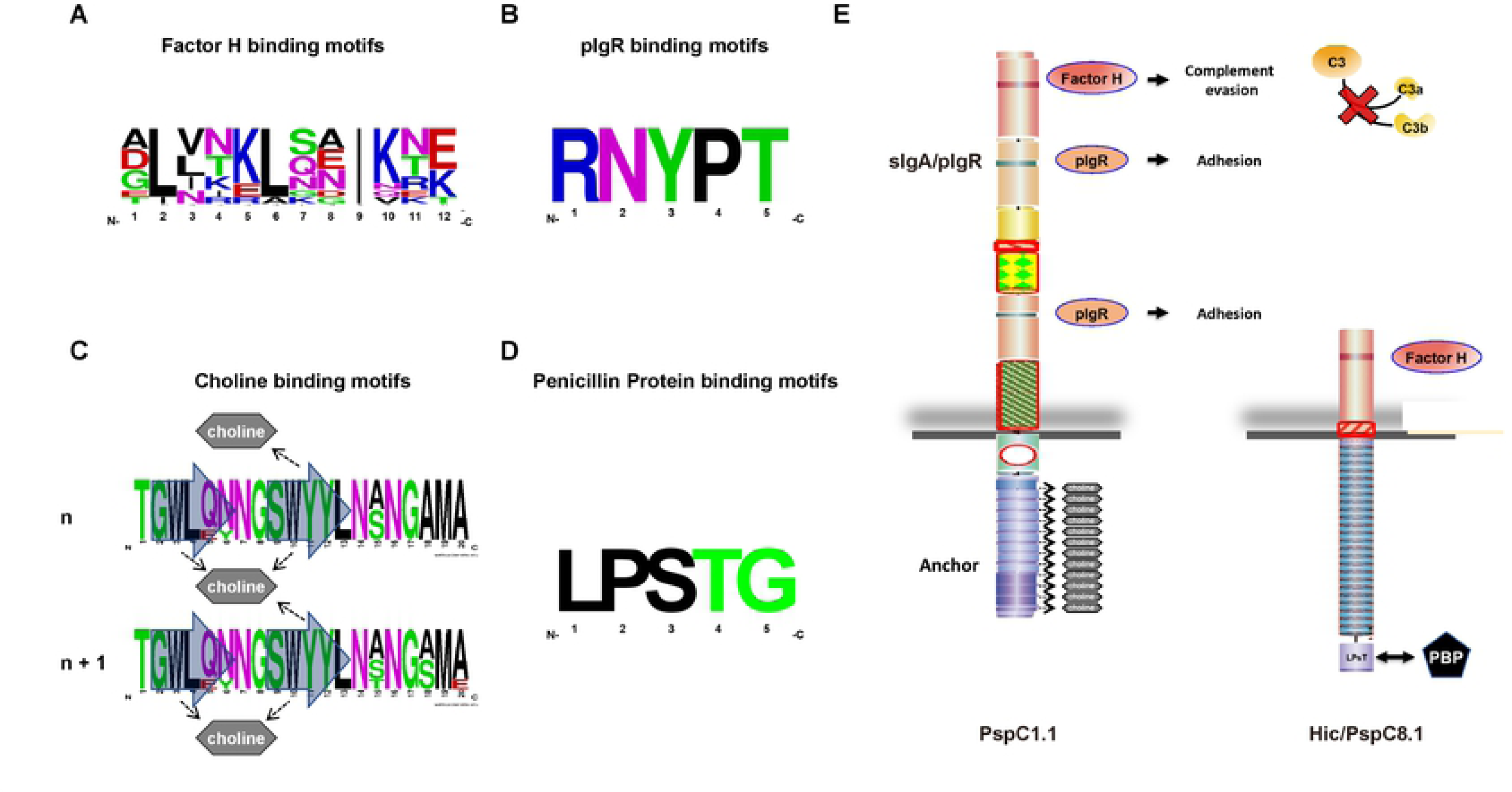
Sequence Variation and Conservation of Binding Domains and Surface Orientation of PspC1.1 and Hic/PspC8.1. **A: Sequence variation of the Factor H binding motif** in the Hypervariable Domains of the six PspC and the five Hic variants. WebLogo was used to evaluate amino acid variation. **B: Sequence conservation in the binding sites for human sIgA/pIgR** in the Repeat Domains I and II. **C: Sequence variation among the Choline-Binding Modules 2 and 3 the PspC variants**. Residues relevant for the contact with choline are indicated by the arrows and include W_3_ and W_10_ of one module, as well as Y_11_ of the following module. WebLogo was used to evaluate sequence variation in the second and third choline-binding modules of the PspC variants. **D: Sequence conservation in C-termini of Hic-type proteins** of the sortase recognition motif LPsTG of covalently anchored proteins. Following sortase cleavage after the T residue and attachment to Penicillin Binding Protein. **E: Structure and proposed orientation of PspC1.1 associated with phosphorylcholine (**PCho)**, and sortase linked Hic/PspC8.1 variant**. The arrangement is based on the concept that PspC1.1 is non-covalently associated to the teichoic acids via its interaction with PCho. In contrast the Hic/PspC8.1 variant is which is covalently linked via the sortase anchor to peptidoglycan Penicillin binding protein (PBP). This attachment and orientation suggests that the Proline-Rich Domains may represent flexible cell wall and capsule spanning segment. Furthermore, the variable length of the C-terminal regions can indicate different types of cell wall attachment, as well as discrete sizes and thickness of the cell wall and the capsule. The grey line represents the bacterial membrane, cell wall and the shaded grey region indicates the position of the capsule. The proposed exterior domains of the PspC and the Hic variant are shown in yellow or red color. The known, mapped binding domains for human plasma regulator Factor H in the Hypervariable Domains (PspC1.1 and Hic/PspC81) and the sIgA or cell surface receptor pIgR in the Repeat Domains I and II (PspC1.1) are indicated by purple and green bars. Attached Factor H mediates complement evasion and blocks complement mediated opsonophagocytosis and release of anaphylatoxins C3a and C5a. SIgA or pIgR bind to two sites in PspC1.1 and avoid opsonization by sIgA or mediate adhesion to human epithelial cells. The binding sites for additional human plasma protein like vitronectin are not mapped so far. The C-terminal region, with a proposed interior location are shown in green, blue or purple color and include the Proline Rich Domains followed by Choline-Binding Module (PspC1.1) or LPsTG mediated anchor (Hic/PspC8.1).

Relationship analysis using a dendrogram identified three subtypes of the hypervariable domains. Subtype A (HVD-A) is present in PspC3.1, PspC5.1, and Hic/PspC11.1 HVD-B is present in the PspC2.2, PspC1.1, and PspC4.2, and HVD-C is present in PspC6.1, Hic/PspC7.1, Hic/PspC10.1, Hic/PspC9.1, and Hic/PspC8.1 (**Supplementary Figure 3C**).

##### Repeat domains

All PspC-type proteins and Hic/PspC7.1 possess approximately 110 aa-long repeat domains (Repeat Domain). Five PspC (i.e. PspC3.1, PspC2.2, PspC6.1, PspC1.1, PspC5.1) possess a second Repeat Domain. The **Repeat Domains** have conserved sequences, they are rich in charged residues, and include conserved RNYPT motifs, which are binding sites for sIgA/pIgR **(Figure 5B, Supplementary Figure 4**). Related repeat domains were identified in PspK from *S. pneumoniae* (H2BJK8) with 55 % homology to Repeat Domain1 and 71.6 % homology to Repeat Domain II, respectively. The solution structure of the Repeat Domain of PspC3.4 from strain TIGR has been solved (40). This domain folds into three antiparallel α-helices, and the YPT residues, representing the core sIgA/pIgR binding motif are positioned in a coiled-coil structured loop, which separates helix 1 and helix 2. This experimentally determined structure actually confirms and validates our *in vitro* structure prediction **(Figure 2A**).

##### Random Coil Domain

Random coil domains are approximately 30 aa-long, show a coiled-coil structure and are relatively conserved in sequence. They are typically positioned downstream of the first Repeat Domain. No homologous were identified in the data bank **(Supplementary Figure 5**).

### New Domains of the N-terminal Region

Sequence stretches in the PspC and Hic variants that did not match known domains of PspC3.1 were also identified. These sequences were used to search for counterparts in other PspC and Hic variants or for homologs in the protein data bank. This procedure identified nine new domains, including one new domain in PspC3.1 and also three new alternates of the Proline Rich Domain.

#### Serine-Rich Elements

Serine-Rich Elements with the overall motif S_n_D/GS_2_ were detected in five PspC and in all Hic/PspC variants. Nine protein variants harbor one, whereas PspC2.2 contains two; and PspC4.2 lacks such an element. These Serine-rich elements share a coiled-coil structure; but differ in position, type, and in sequence. Ser-rich elements following the Hypervariable Domain (PspC2.2, Hic/PspC7.1, Hic/PspC9.1, Hic/PspC8.1) or the unique Hic/PspC11.1 domain have the consensus S_n_D/GS_2_ and are up to 24 aa long. The segments following the Random Coil Domain (PspC3.1, PspC2.2, PspC6.1, PspC1.1, PspC5.1, Hic/PspC10.1) have related S_2_DS_2_, units, which can be up to 18 aa long. The domain of Hic/PspC10.1 shows a variation to these common features **(Supplementary Figure 6A**). The biological role(s) of these elements are as yet unknown. In engineered proteins, related poly-serine-rich elements are integrated as flexible linkers that separate functional, individually folding domains (41). Interestingly the TKPET motif at the end of segments following the Hypervariable domains are highly related to the first seven residue long units found in Proline Rich Domains III and IV (see below).

### Random Coil Extension Domains

Two separate new domains were identified in four proteins, which are positioned downstream of the Random Coil Domain-S_2_DS_2_ combination.

#### Random Coil Extension Domain 1

Two proteins, PspC1.1 and PspC5.1, contain a new domain following the Random Coil Domain-S**_2_**DS_2_ combination. These 83 aa long domains share almost identical sequences.

This domain includes several charged residues, and shares homology with domains in other proteins. Proteins containing such RICH type domains including secreted proteins such as PspC Q9KK19, SpsA O33742 and IgA Fc receptor binding protein P27951 from *Streptococcus agalactiae*. The domain is predicted to be involved in bacterial adherence or cell wall binding (42).

#### Random Coil Extension Domain 2

PspC4.2 and Hic/PspC10.1 have different 114 (PspC4.2) or 126 aa-long (Hic/PspC10.1) segments following Random Coil Domain- S_2_DS_2_. These elements show moderate sequence homology among each other. The 126 aa domain of Hic/PspC10.1 harbors a N-terminal 37 aa extension, with the remaining portion being conserved in the PspC4.2 domain. The biological role of this unique segment is unclear. In PspC4.2 the region includes a long α-helical stretch and is followed by a ca 30 residue long coiled-coil stretch.

#### PspA-Like Domain

PspC1.1 and PspC5.1 have related, new domains following Repeat Domain II. These 131 or 130 aa-long domains are rich in charged residues, and exhibit 84.5% sequence homology with the A*/B element of *S. pneumoniae* PspA from strain DBL6A, which includes a lactoferrin-binding region (43, 44). These data suggest that the newly identified domains in PspC1.1 and PspC5.1 bind lactoferrin (45, 46).

#### PspC4.2 Specific Element

Domain pattern analysis identified an element in PspC4.2 which is positioned between the Hypervariable Domain and the Random Coil Domain. This 33 aa-long α-helical structured segment, lacks homology to other proteins in the databank role, thus its role remains unclear.

#### Repeat Type Domain

PspC4.2, Hic/PspC7.1, and Hic/PspC10.1 share related 92, 82, or 68 aa-long domains, which are distantly related (41.6% homology) to the Repeat Domains. These new Repeat Type Domain with a mostly α-helical structure intriguingly, lack an RNYPT binding motif and seem to be specific for PspC and Hic proteins.

### A New Two Segmented Domain

A new two-domain segment was identified in PspC4.2 and the three Hic proteins, Hic/PspC7.1, Hic/PspC10.1, Hic/PspC9.1.

#### The upstream domain

The 24, 36, 40 or 37 aa-long upstream sections are rich in proline residues (Supplementary Figure 14), have a predicted coiled structure, and due to their location in the N-terminal region are termed ***Extracellular Proline Rich Segments***. The high Proline content may suggest a function as chain breaker (47). These External Proline Rich Domains lack homology to other bacterial proteins, and thus seem unique for PspC proteins.

The downstream units show homology to the Fc binding part of protein C from S. agalactiae The 89 or 78 aa long elements are rich in charged residues, lack proline residues, and have an α-helical structure. A blast search revealed 51.1% identity to a segment within the trypsin sensitive beta-antigen of *Streptococcus agalactiae* (strain P27951/Uniprot). This protein binds the Fc region of human IgA likely via two stretches (48). This sequence stretch is found in several bacterial immune evasion proteins. One group of Gram-positive bacteria share in their signal peptides a YSIRK motif. Also based on the many charged residues this domain (pfam05062) is also named RICH (Rich In Charged residues) and is identified in other secreted proteins of *S. pneumoniae* proteins including SpsA and the Fc binding part of human IgA from *Streptococcus agalactiae*. The function is proposed in bacterial adherence or cell wall binding.

#### Hic/PspC11 Specific Element

Hic/PspC11.1 contains a unique 102 aa-long α- helical structured domain, which follows the Hypervariable Domain. Related segments were identified in most Hic/PspC11 variants, however, not in other pneumococcal or bacterial proteins Thus far, the function of this domain is unknown..

### Domain Composition of the C-terminal region

The C-terminal regions of the analyzed PspC and Hic proteins are relatively conserved in length (ranging from 237 aa (PspC5.1) to 348 aa (Hic/PspC8.1) and each protein combines a modular Proline-Rich Domain with either the PspC specific Choline-Binding Domain or the Hic characteristic LPsTG anchor (47–50). A general pattern is emerging: PspC proteins link shorter Proline-Rich Domains (57 to 77 aa) to longer Choline-Binding Domains (179 to 219 aa), while Hic proteins combine longer, Prolin-Rich Domains (186 to 286 aa) with shorter LPsTG anchors (50 to 62 aa).

#### Proline-Rich Domains

Proline-Rich Domains have a modular structure and connect the N-terminal region with the cell wall anchor. The proposed role as a bacterial cell wall-spanning domain is consistent with the position prior to the anchor (49, 50). Our in silico analysis identified a modular composition and further distinct proline-rich domains, which differ in length (57 to 286 aa), type, module composition, and sequence.

#### Proline-Rich Domain I

Five PspC variants have highly related 59 to 77 aa long domains, forming either three-segmented (PspC3.1, PspC6.1, PspC2.2), or two-segmented domains (PspC1.1, PspC5.1)(**Supplementary Figure 7A**). The first N-terminal segments have Proline dominated PAPA- and PAPAP motifs, and can be up to 46 aa long. The C-terminal segments include PAPAP or PAPTP-forming motifs, are up to 19 aa long, and have a coiled-coil structure. The middle segment found only in the three segmented domains is conserved in length (23 aa), in sequence, exhibits characteristic flanking Q-residues, and is rich in charged residues. Interestingly this segment has a predicted α-helical structure and lacks Prolines. Such Proline-Rich segments are also found in PspA (50, 51).

#### Proline-rich domain II

PspC4.2 has a unique 57 aa-long Proline-Rich Domain. This new domain includes 19 Prolines and has an internal repeated segment with the sequence TPQV**P**K**P**EA**P**K. To date, this new domain has been identified only in PspC proteins)(**Supplementary Figure 7B**).

#### Proline-rich domain III

Hic/PspC7.1 harbors a unique 186 aa-long Proline-Rich Domain which includes an N-terminal 7 aa element followed by five almost identical 31 aa long repeats (KK**P**SA**P**K**P(**G/D)MQ**P**S**P**Q**P**EGKK**P**SV**P**AQ**P**GTED). Each repeat has **nine** Prolines and two KKPS(A/V)P motifs (denoted by white letters). The 31 aa repeats are followed by a truncated 24 aa-long repeat element (**Supplementary Figure 7C, D**).

#### Proline-rich domain IV

Four Hic variants harbor 247 to 286 aa long, Proline-Rich Domains containing 23, 19, or 26 modules. The modules vary in type and sequence, including multiple 11 aa modular repeats, which are followed by one truncated repeat, and a 16 aa long extension (**Supplementary Figure 8A,8B,8C**). Hic/PspC10.1 and Hic/PspC9.1 contain 14 and 16 modular units, respectively with the sequence (L/P)E**K**PKPEVKP**Q**. Hic/PspC8.1 and Hic/PspC11.1 contain 23 copies of (L/P)E**T**PKPEVKP**E** elements (variant residues in white letters on black background). They are followed by one shortened module and have distal nearly identical 16 aa-long C-terminal units, which at position 15 show a **T/P** variation)(**Supplementary Figure 8D, 8E, 8F**).

#### Cell wall attachment

PspC proteins use longer modular Choline-Binding Domains for cell wall attachment and by contrast, Hic proteins have shorter, 50–62 aa-long anchors that include a sortase-dependent LPsTG motif (53, 54).

### PspC-type protein variants possess choline-binding anchors

PspC type, variants have modular C-terminal Choline-Binding Domains and their length ranges from 178 (PspC5.1) to 248 aa (PspC1.1). Most modules are 20 aa and apparently two units can attach to one choline component (52). Related Choline-Binding Domains are found in up to 15 other *S. pneumoniae* proteins, including the immune evasion protein PspA, the autolysins LytA, LytB LytC, and CbpL (52). In the literature these modular composed Choline-Binding Domains are sometimes termed choline- binding modules. However given the domain composition of full length PspC and Hic variants, we prefer to term such smaller, repetitively assembled subunits as modules. Apparently both PspC and Hic variants use modular composited domains within their C-terminal putatively interior regions.

#### Hic variants have C-terminal sortase signals

The five analyzed Hic variants share C-terminal 50–62 aa anchors which display a specific pentapeptide LPsTG motifs. The transpeptidase, sortase A cleaves within this conserved motif between Thr and Gly, and subsequently the protein is covalently linked via the Thr to lipid II (P3 precursor) and a penicillin binding protein (55, 56)(**Figure 5E**). The domain distribution of related PspC and Hic variants show differences which are indicative for extra and intracellular positions, and reveal diverse structural composition among PspC and Hic variant proteins.

## Discussion

This analysis of domains within the six PspC and five Hic Hic variants identified 13 N-terminal and three C-terminal domains, including the seven known and nine new domains, and furthermore recognized three new alternates of the Proline-Rich Domain. The mature PspC and the Hic proteins are heterogeneous proteins. They generate intriguing diversity by combining different domains, by varying domain types, and by assembling domains in different numbers. Domain variability is increased by assembling variant modular elements and by sequence variations. This reflects antigenic variation, functional specialization and furthermore the different anchors are indicative for a different surface distribution (17, 18). Three domains, the Signal peptides, the Hypervariable Domains and Proline-Rich Domains are found in all analyzed variants (Table I). Eleven domains are found in several (but not all) variants, and two domains are unique to single proteins. This extensive domain characterization shows differences between the analyzed PspC and Hic variants, reveals variable domain assembly features, and shows a different composition of the N and C-terminal regions.

### Variability among PspC and Hic-variants

PspC, and Hic-type S pneumonia show related domains in their N-terminal regions, but differ more in their C-terminal regions. Also the proteins have different C-terminal anchors. PspC proteins with the Choline-Binding Domains contact multiple choline-moieties in a non-covalent manner. In contrast the LPsTG anchors attach the proteins covalently to the peptidoglycan (54). The type of C-terminal anchor not only influences cell wall attachment, but length and composition of the Proline-Rich Domains and furthermore seems to influence selection, composition, and number of the N-terminal domains.

These different domain pattern distributions are indicative for distinct surface positioning and different roles in immune evasion.

#### Variability of N vs C-terminal regions

Broadly speaking, each PspC and Hic protein has two major parts: the N-terminal, outside presented region, which includes immune evasion and adhesion domains, and the C-terminal anchoring region.

The **N-terminal** regions of the analyzed PspC and Hic proteins vary in length, and domain number ranging from 155 aa with two domains (Hic/PspC8.1) to 610 aa with eight domains (PspC4.2). These regions share structural features, including long α-helical structures, and a high proportion of charged residues. The Hypervariable Regions are located most distant from the cell surface and show the highest degree of variation. This diversity reflects differences in immune control and antigenic variability, which is relevant for evading immune recognition by antibodies. Six of the N-terminal domains are unique to PspC and Hic variants, others like the PspA Related Domain and the region with homology to the IgA binding β antigen are found in other pneumococcal or bacterial immune evasion proteins.

**The C-terminal** regions are distinct from the N-terminal regions. They are more conserved in length, ranging from 236 aa (PspC5.1) to 348 aa (Hic/PspC8.1), have more polar and amphipathic residues and PspC proteins have also more Tyr residues. The Proline-Rich Domains, preceding the PspC and Hic-specific anchors, show a modular composition, have mostly coiled-coil structures and differ in length. Proline-Rich Domains of PspC proteins are shorter than those of Hic proteins. Given the proposed location at the interface between cell wall and capsule, such diversity could reflect different binding dynamics, strength of cell wall integration, or morphological differences due to capsule thickness (54–59). Similarly the anchor domains in the C-terminus differ in length, composition, and type of cell wall integration.

#### Protein orientation, and cell wall integration

PspC and Hic are membrane integrated, surface proteins and we are understanding now which parts of the proteins have exterior or interior location. The N-terminal region, by extending from the capsule, is exposed to the outside world and can interact with human proteins (Table II). The C-terminal region includes a capsule spanning segment and an internal cell wall anchor.

**Table II:** Domains Identified in the evaluated PspC and Hic variants. The domains are listed from N-terminal to the C-terminal region, and known as well as new domain and domain alternates are presented. Also domains which are specific for PspC/Hic variants are shown, as well as domains which are shared and found in other *S. pneumoniae* proteins and in other bacterial proteins. RDSP signal peptide; HVD hypervariable Domain; RD Repeat Domain; RCD Random Coil Domain; SnD/GS2 Serine Rich segment; RCE Random Coil Extension; R-type repeat related Domain; EPRD Extracellular Proline Rich Domain; VS Variant specific; IgA IgA Binding Domain, PRD Proline-Rich Domain, CBP Choline-Binding Domain.

| Host Regulator | Function | Binding Site |
| --- | --- | --- |
| Factor H | Complement Regulation | HVD |
| sIgA/plgR | Adhesion | Repeat Domains |
| C3 | C3 Inactivation | Not mapped |
| C4BP | CP Inhibition | Not mapped |
| Plasminogen | Proenzyme; plasmin cleaves<br>inactivates C3, C3b and fibrin | Not mapped |
| Thrombospondin-1 | adhesive glycoprotein, cell-<br>cell and cell-matrix interaction | Not mapped |
| Vitronectin | Complement control &<br>adhesion | Not mapped |
| Lactoferrin | Fe metabolism | Proposed by homology |
| IgA | IgA Inactivation? | Proposed by homology |

Cell wall attachment via the C-terminal anchor orients the N-terminus to the outside to allow interaction with host plasma proteins and cell receptors. An illustration of the orientation, spatial organization of one PspC and one Hic variant including mapped binding sits for human plasma regulators is presented in **Figure 5E**. PspC1.1 representing an eight domain choline-attached variant and the short four domain Hic variant (Hic/PspC8.1) show different compositions both in the proposed extra and intracellular regions. Due to variable length the N-terminal regions extend to different distances from the surface. In a linear model, for example, Factor H, when bound via the hypervariable domain inhibits C3b formation and assists in C3b inactivation remote from the bacterial surface. Similarly, the Proline-Rich Domains, due to their variable length and composition, and the specific anchors can integrate the proteins in the cell wall envelope with different depth and strength.

#### Tactical positioning and immune evasion

The two distinct anchors, show different structures, mainly β-sheet composed Choline-Binding Domains vs coiled-coiled and α-helical structured LPSTG anchors. This not only mediates non-covalent vs. covalent attachment, but is also indicative of a more flexible vs. fixed cell wall attachment, for a different surface distribution and probably also exposure to the host. Indeed, for *S. pneumoniae* strain BNH418, a different spatial localization of a PspC and a Hic variant was shown by super resolution microscopy (60). The PspC- protein, with the Choline-Binding Domain localized to the division septum and bound Factor H as a result controlled C3b opsonization. In contrast, the LPsTG anchored Hic protein was localized to the bacterial poles. Such distinct surface localization can apparently influence the site of complement control, adhesion to host cells and can reflect a tactical positioning of these important immune evasion proteins. This structure and sequence based analyses suggests that differences between PspC and Hic variants extend beyond cell wall anchors to other domains and to domain pattern usage.

When comparing prevalence and distribution of PspC vs. Hic variants among 349 clinical *S. pneumoniae* isolates derived from adult patients with invasive pneumococcal disease, 298 isolates (85.4%) encoded a PspC-variant, 22 strains (6.3%) a Hic-variant, 19 isolates (5.4%) had a *pspC* and *hic* genes and only 10 isolates (2.9%) had neither *pspC* nor *hic* genes (61). In addition, invasive, PspC expressing strains bound more Factor H, and Factor H binding and immune control was more effective in encapsulated as compared to unencapsulated strains. Similarly the PspC variants (i.e. PspC2 and PspC6) were more efficient in Factor H binding and complement inhibition on the bacterial surface as compared to Hic variants (Hic/Pspc9 and Hic/PspC11) (62, 63).

#### Conclusions and Perspectives

Evaluating the domain composition of selected PspC and Hic variants and an in-depth characterization of the domain composition resulted in a better understanding of the structure and role of these pneumococcal virulence determinants in immune evasion. Our approach identified further differences between PspC and Hic proteins, which are beyond their distinct membrane anchors. Such knowledge allows a comparison of full-length proteins based on domain patterns, numbers and composition and can result in a better comparison between PspC and Hic proteins. Similarly, individual domains can be compared based on structure, modular composition and sequence.

Analyzing the additional >60,000 PspC and Hic proteins deposited in the NCBI protein data bank or gene products from new clinical isolates, will likely identify additional variants, new domains, novel domain combinations, and also new subdomains. Understanding the composition of these diverse pneumococcal virulence factors will better explain their role in immune evasion, provide important information for molecular strain typing, and for vaccine design. Last but not least this may also allow a correlation between PspC or Hic type variants with invasive pneumococcal infections and with clinical outcome e.g. of young patients with Pneumococcal Hemolytic Uremic Syndrome.

## Materials and method

### Selection of PspC and Hic variant proteins

Each of the selected six PspC and five Hic proteins represents one of the two cluster as initially defined by Ianelli *et al.* (38). The sequences were derived from the NCBI protein site (status: **Feb./2018**). The general PspC /Hic designation is based on the definition by Iannelli,et al. (38). The protein names, corresponding bacterial strain, protein size, GenBank Accession number and protein ID are shown in (**Supplementary Table I).**

### Secondary structure evaluation

The structure (α−helical, coiled-coil and β−sheet) of each selected PspC and Hic protein was evaluated using the program presented via the RaptorX server (http://raptorx.uchicago.edu/). The PspC3.1 shows best matched template is ***2vyuA*** with p-value:3.39e-10 and secondary structure: 42% (α−helical, 43% coiled-coil and, 14% β−sheets. The other ten PspC Hic variants were evaluated in the same manner, and showed a similar secondary structure composition (Supplementary Figures 1- 10). Each of the six PpsC variants matched best to the same template: 2vyuA, and the five Hic variants (Hic/PspC7.1, Hic/PspC8.1, Hic/PspC9.1, Hic/PspC10.1, Hic/PspC11.1) matched to templates (1w9rA, 4k12B, 2m6uA, 6iaA, 2m6uA, respectively). Three-class secondary structure prediction results are shown in the form of histograms which were constructed using ggplot2 from the R/Bioconductor.

### Phylogenetic analysis

The PspC and Hic protein sequences and amino acid composition were evaluated using MEGA7 (www.megasoftware.net). There were a total of 976 positions in the final dataset. Evolutionary analyses were conducted in MEGA7 (62. Kumar S., 2016). The CLUSTALW program and the BLOSUM amino acid matrix was used to compare the allelic variants of PspC, following which phylograms were generated using the Neighbor-Joining method (Bootstrap value:100). Each domain’s phylogram was generated by the same method described for the full-length protein sequences. Phylogenetic trees are modified in MEGA7.

### Domain blast analysis

The software BLASTp was used to conduct homology searches of the GenBank database available at the National Center for Biotechnology Information website (http://www.ncbi.nlm.nih.gov/). Furthermore the software BLAST targeting database UnipRotKB reference proteomes plus Swiss-Prot was used to find regions of local similarity between sequences (https://www.uniprot.org/blast/). All the domains in this work have been done a blast.

## Acknowledgments

The work of the authors is supported by the Collaborative Research Center, FungiNet (projects C6 (PFZ) and C4 (CS)) Deutsche Forschungsgemeinschaft (DFG). SL acknowledges a fellowship from the German Academic Exchange Service (DAAD) and from the ILRS, International Leibniz Research School for Biomolecular Interaction, Jena, Germany. Alfredo Sahagún-Ruiz was funded by a scholarship from PASPA-DGAPA, National Autonomous University of Mexico (UNAM), and from Mexican National Science and Technology Council (CONACYT) for a sabbatical stay at Department of Infection Biology, Leibniz Institute for Natural Product Research and Infection Biology - Hans Knöll Institute, Jena, Germany. SH received funding by the Deutsche Forschungsgemeinschaft DFG HA 3125/5-2.

## Supplementary Figure Legends

**Supplementary Figure 1: Structural composition of PspC variants with choline binding anchors. (A)** The structure of Psp2.2 was evaluated *in silico*. The N- terminus shows a long stretch composed mainly of α-helices (red columns) (aa 1- 440), being followed by a 74 aa long coiled-coil structured segment (grey area) and by a 179 aa long segment with β−sheet folds (blue columns). The signal peptide (aa 1-37) which is cleaved upon processing is shown by the grey background and blue lines. The vertical grey bar separating the N-terminal α helical region and the C- terminal coiled coil structured region may represent the position of the bacterial cell wall. **(B) Structural composition of PspC2.2.** The structure of Psp2.2 was evaluated *in silico*. The N-terminus shows a long stretch composed mainly of α- helices (red columns) (aa 1-405), being followed by a 77 aa long coiled-coil structured segment (grey area) and by a 199 aa long segment with β−sheet folds (blue columns). The signal peptide (aa 1-37) which is cleaved upon processing is shown by the grey background and blue lines. The vertical grey bar separating the N- terminal α helical region and the C-terminal coiled coil structured region may represent the position of the bacterial cell wall. **(C) Structural composition of PspC1.1.** The structure of PspC1.1 was evaluated *in silico*. The N-terminus shows a long stretch composed mainly of α-helices (red columns) (aa 1-626), being followed by a 64 aa long coiled-coil structured segment (grey area) and by a 248 aa long segment with β−sheet folds (blue columns). The signal peptide (aa 1-37) which is cleaved upon processing is shown by the grey background and blue lines. The vertical grey bar separating the N-terminal α helical region and the C-terminal coiled coil structured region may represent the position of the bacterial cell wall. **(D) Structural composition of PspC5.1.** The structure of Psp5.1 was evaluated *in silico*. The N-terminus shows a long stretch composed mainly of α-helices (red columns) (aa 1-632), being followed by a 59 aa long coiled-coil structured segment (grey area) and by a 178 aa long segment with β−sheet folds (blue columns). The signal peptide (aa 1-37), which is cleaved upon processing is shown by the grey background and blue lines. The vertical grey bar separating the N-terminal α−helical region and the C-terminal coiled-coil structured region may represent the position of the bacterial cell wall. **(E) Structural composition of PspC4.2.** The structure of Psp4.2 was evaluated *in silico*. The N-terminus shows a long stretch composed mainly of α-helices (red columns) (aa 1-610), being followed by a 57 aa long coiled-coil structured segment (grey area) and by a 199 aa long segment with β−sheet folds (blue columns). The signal peptide (aa 1-37) which is cleaved upon processing is shown by the grey background and blue lines. The vertical grey bar separating the N- terminal α-helical region and the C-terminal coiled-coil structured region may represent the position of the bacterial cell wall.

**Supplementary Figure 2: Structural composition of Hic type variants with LPsTG anchors. (A) Structural composition of Hic/PspC7.1.** The structure of HIC/Psp7.1 was evaluated *in silico*. The N-terminus shows a long stretch composed mainly of α-helices (red columns) (aa 1-533), being followed by a 186 aa long coiled-coil structured segment (grey area) and by a 50 aa long segment with an LPsTG motif. This segment has mostly α−helical structure. The signal peptide (aa 1-37) which is cleaved upon processing, is shown by the grey background and blue lines. The vertical grey bar separating the N-terminal α-helical region and the C-terminal mostly coiled-coil structured region may represent the position of the bacterial cell wall. **(B) Structural composition of Hic/PspC10.1.**The structure of HIC/Psp10.1 was evaluated *in silico*. The N-terminus shows a long stretch composed mainly of α- helices (red columns) (aa 1-502), being followed by a 204 aa long coiled-coil structured segment (grey area) and by a 57 aa long segment with an LPsTG motif. This segment has preceding the motif a coiled coil and α−helical structure following the motif. The signal peptide (aa 1-37) which is cleaved upon processing, is shown by the grey background and blue lines. The vertical grey bar separating the N- terminal α-helical region and the C-terminal mostly coiled-coil structured region may represent the position of the bacterial cell wall. **(C) Structural composition of Hic/PspC9.1.** The structure of HIC/Psp9.1 was evaluated *in silico*. The N-terminus shows a long stretch composed mainly of α-helices (red columns) (aa 1-279), being followed by a 247 aa long coiled-coil structured segment (grey area) and by a 57 aa long segment with an LPsTG motif. This segment has preceding the motif a coiled coil and α−helical structure following the motif. The signal peptide (aa 1-37) which is cleaved upon processing is shown by the grey background and blue lines. The vertical grey bar separating the N-terminal α−helical region and the C-terminal mostly coiled-coil structured region may represent the position of the bacterial cell wall. **(D) Structural composition of Hic/PspC8.1.** The structure of HIC/Psp8.1 was evaluated *in silico*. The N-terminus shows a long stretch composed mainly of α- helices (red columns) (aa 1-155), being followed by a 286 aa long coiled-coil structured segment (grey area) and by a 62 aa long segment with an LPsTG motif. This segment has preceding the motif a coiled coil and α−helical structure following the motif. The signal peptide (aa 1-37) which is cleaved upon processing is shown by the grey background and blue lines. The vertical grey bar separating the N-terminal α-helical region and the C-terminal mostly coiled-coil structured region may represent the position of the bacterial cell wall. **(E) Structural composition of Hic/PspC11.1.** The structure of HIC/Psp11.1 was evaluated *in silico*. The N-terminus shows a long stretch composed mainly of α-helices (red columns) (aa 1-264), being followed by a 286 aa long coiled-coil structured segment (grey area) and by a 62 aa long segment with an LPsTG motif. This segment has preceding the motif a coiled coil and α−helical structure following the motif. The signal peptide (aa 1-37) which is cleaved upon processing, is shown by the grey background and blue lines. The vertical grey bar separating the N-terminal α-helical region and the C-terminal mostly coiled coil structured region may represent the position of the bacterial cell wall.

**Supplementary Figure 3: Amino acid sequences of Signal Peptides and of the**

**Hypervariable Domains.**

**A: Sequence Conversation of the** Signal Peptide. Sequences of the N-terminal region of the six PspC and the five Hic/PspC variants are shown. Conserved residues are shown with a black background; residues which are present in most proteins are shown on grey background. Positively charged residues are shown in blue, and negatively charged residues in red characters. **B: Sequences of the Hypervariable Domains** of the six PspC and the five Hic variants. The Factor H binding sites which has been mapped for PspC3.1 is shown with green background. The hypervariable domains can be separated into three major groups, termed HVD- A, HVD-B and HVD-C.

**Supplementary Figure 4: Sequence Conservation in Repeat Domains I and II**

**A:** Sequences of the N-terminal region of the Repeat Domains of six PspC- and the HIC/PspC variant. See legend to Supplementary Figure 11 for explanation. The conserved binding domain for sIgA is shown with yellow background. **B:** Alignment of Repeat Domain II.

**Supplementary Figure 5: Conserved Residues of the Random Coil Domains and S_n_D/GS_2_ Domains.**

**A**: Sequences of the Random Coil Domain following the first Repeat Domain are shown. See legend to Supplementary Figure 3 for explanation. **B**: **Conserved Residues in S_n_D/GS_2_ Domains.** The upper panels show the segments that follow the Hypervariable regions and the lower panel the Proteins and segments that follow the random Coil domain.

Supplementary Figure 6: **Conserved Amino Acid Residues of the New N- terminal Domains.**

**A:** Sequence Homology of the New Random Coil Extension region I**; B:** Sequence Homology of the New Random Coil Extension region II **C:** Sequence Homology of the **Random Coil Domain; D: Homology of PspA like Domains.** The bottom row shows the sequence of PspA from strain EF6769 with includes a lactoferrin binding domain. E: **Homology of Repeat Relate** The bottom row shows the sequence of PspA from *S. agalactiae.* **F: Sequence of the PspC4.2 specific Element G: Sequence of the Hic/PspC11.1 Specific Domain. General:** Conserved residues are shown with a black background; residues which are present in most proteins are shown on grey background. Positively charged residues are shown in blue, and negatively charged residues in red characters.

Supplementary Figure 7: **Conserved Residues of the C-terminal Proline-Rich Domain I to Proline-Rich Domain III.**

**A: Proline-Rich Domain I.** Proline Rich Doman I is used by five PspC proteins, PspC3.1, PspC2.2, PspC6.1 PspC1.1, PspC5.1. This domain has three major regions. The first and third domains have conserved Pro residues and the motifs PAPAP and PAPAT are found in most domains. The C-terminal part is also rich in Pro residues. The middle segment, element II which is represented by flanking Q- residues, is found in PspC1.1, PspC2.2, and PspC6.1 but is absent in PspC1.1 and PspC5.1. **B: Proline-Rich Domain II.** PspC4.2 uses a separate repeat domain which has an 18-residue element duplicated. **C: Proline-Rich Domain III.** Hic/PspC7.2 has a Proline Rich Domain composed of six segments. The first segment is seven residues long, segments 2-5 include duplicated 31 residue long regions and the most C-terminal segment represents is a truncated 24 aa long version of these repeat units. **D:** The conserved and variant residues were identified by WEBLOGO.

**Supplementary Figure 8: Conserved Amino Acid Residues of the C-terminal Proline-Rich Domains IV.**

**Proline-Rich Domain IV of Hic/PspC9.1 and Hic/pspC10.1.** Variants of the Proline- Rich Doman IV is used by Hic/PspC9.1(A) and Hic/PspC10.1 (B). Both domains use a six residue long segment as first unit, which are followed, by four (Hic/PspC9.1) or two (Hic/PspC10.1) 11 amino acid long segments. Next segments 6-21 (Hic/PspC9.1) or segments 4-17 (HIC/PspC10.1) are 11 residues long and highly related to each other. The most C-terminal segment is 22 residues long unit. **C:** Sequence homology alignment of the conserved 11 residues long elements of PspC9.1 and Hic/PspC10.1. **C**: conserved and variant residues in Hic/PspC9.1 and Hic/PspC10.1 The Variation occurs at position 1 of the repeat units. WEBLOGO was used for alignment. **Proline-Rich Domain VI. Highly related variant of the** Proline Rich Doman VI are used by Hic/PspC8.1 **(D**) and Hic/PspC11.1 (**E**). Both domains use a seven residue long segment as first unit. Units 1-24 are highly related almost conserved 11 residue long repeated units. A variation occurs at position 1 of the repeat units. **F:** Homology alignment of the conserved 11 residues long elements of Hic/PspC8.1 and Hic/PspC11.1 using WEBLOGO. A variation occurs at position 1 of the repeat units.

**Supplementary Figure 9: Conserved amino acid residues of the C-terminal anchor sequences of the PspC and Hic variants.**

**A:** Sequence Homology among the modules of the Choline Binding Domains in the six PspC variants, i.e. PspC3.1, PspC2.2, PspC6.1, PspC1.1, PspC5.1, PspC4.2 are shown. Based on sequence comparison the first modules, have a more different amino acid structure and show conserved sequence pattern which identified these domains as the first positon in a choline binding domain.. The following modules form a middle segment, where the modules are also relatively conserved to each other in sequence. This middle segment shows variation in number of modules, ranging from five to eight. Similarly the three most C-terminal modules of each domain show position specific features and they are conserved among the six PspC variants. These modules are termed third to last, second to last and last most C terminal module.**B:** Sequence alignment of the LPsTG anchor of the five HIC variants. The **LPsTG** motif which is relevant for sortase anchoring is shown in white letters on red background.

Supplementary Table I: **Proteins representing the specific clusters which were selected for the detailed Sequence and structural Analyses.** Protein names, strain origin, length in aa, as well as GenBank and Protein Accession numbers are presented.

